# Spatial Transcriptomic Profiling Reveals Microenvironment-Dependent Immune Signatures in a Lyme Arthritis model

**DOI:** 10.64898/2026.06.25.734510

**Authors:** Kaitlyn Gura, Mollie Hostetter, Veena Potluri, Michael Hill, Sarah Johnson, Yuelin Zhong, Eleanor Astley, Tanja Petnicki-Ocwieja, Suba Nookala, Catherine Brissette, Archana Dhasarathy

## Abstract

Lyme arthritis, a manifestation of Lyme disease, is triggered by the spirochetal bacterium *Borrelia burgdorferi* (*Bb*), which is transmitted through the bite of the *Ixodes* tick. Although multiple studies have been conducted on the complex host immune response in Lyme arthritis, the spatial gene expression environment in the joint tissue remains unexplored. Here, we applied spatial transcriptomics to ankle joints of C3H mice infected with *Bb*, profiling tissues at peak inflammation (two weeks post infection) and after antibiotics (four weeks post-infection) during inflammation resolution. Analysis revealed spatially restricted signatures: pro-inflammatory responses dominated synovial and fibroblast populations two weeks post-infection, with elevated levels of *Vimentin* and I-E*^k^* gene - and *Vimentin* protein - expression localized to these regions. By four weeks post-infection during the inflammation resolution phase, levels of *Vimentin* and *I-E^k^* related gene and protein expression were reduced. Further, we noted an increase in the CD54^+^ and CD106^+^ double-positive population in infected mice joints compared to the vehicle treated controls. Notably, fibroblasts and synoviocytes in the medial joint regions adopted immune-like phenotypes during peak inflammation, while the same cell types in the exterior humeroradial joint displayed a more infection-resilient phenotype. These spatially resolved maps demonstrate that joint microenvironments play a crucial role in pathogenesis, offering unique insights into Lyme arthritis pathology.

## Introduction

Lyme disease, caused by the spirochete *Borrelia burgdorferi (Bb)*, represents the most common vector-borne illness in North America and Europe, with over 476,000 annual cases estimated annually in the United States alone. Lyme arthritis (LA) develops in approximately 60% of untreated Lyme disease patients and persists despite antibiotic therapy in about 10% of cases, a phenomenon known as post-treatment LA [1].

Multiple cell types are found in the joint, including synoviocytes, macrophages, T-cells, neutrophils and endothelial cells, of which fibroblast-like synoviocytes constitute the primary cell type [2–4]. The interplay between these cell types and signaling pathways are critical for development of arthritis following *Borrelia* infection [5]. The dysregulation of these interactions and /or immune pathways likely underlie the development of Lyme arthritis. Prior transcriptomic analyses of LA have provided valuable insights into these dysregulated immune pathways, despite their use of bulk RNA sequencing, which effectively homogenizes tissue samples, thereby averaging gene expression across diverse cell types and spatial contexts. Additionally, cell-type specific, rare immune transcripts were often masked by bulk analyses. The advent of more recent methods such as single-cell RNA sequencing and spatial transcriptomics has helped resolve transcription changes in the context of spatial and cellular neighborhoods, leading to better understanding of the tissue microenvironments driving pathology.

In particular, spatial transcriptomics has helped characterize the spatial relationships between infiltrating cells and resident cells, thereby enabling genome-wide gene expression profiling while preserving native tissue architecture [6]. This technology uncovers cellular organization, transcriptional niches, and patterns of immune infiltration, proving especially valuable for dissecting complex inflammatory processes in joint tissues. For instance, spatial transcriptomics of human rheumatoid arthritis synovium identified distinct inflammatory microenvironments around tertiary lymphoid organs, with co-localized B cells and other leukocytes driving chemotaxis and humoral responses not visible in homogenized samples [7]. In another study, spatial transcriptomics in osteoarthritis synovium revealed disrupted immune homeostasis, mapping 27 cell subclasses into lining/sub lining domains and identifying CD69 attractors for myeloid and lymphoid infiltration. To our knowledge, this technology has yet to be applied to Lyme arthritis models.

Here, we present the first spatial transcriptomic analysis of *Bb* infected C3H murine joints, enabling characterization of spatial gene expression patterns and cellular organization during peak inflammation (2 weeks post-infection) and following antibiotic-mediated resolution (4 weeks post-infection). We used C3H mice, which are highly susceptible to *Bb* infection and develop robust inflammation that recapitulates key pathological features of arthritis, including synovial hyperplasia, immune cell infiltration, and cartilage damage. Importantly, C3H mice also exhibit Lyme arthritis that presents similarly to subacute Lyme arthritis in humans, making them ideal for studying persistent inflammation [8]. Our findings provide unique insights into the temporal and regional complexity of Lyme arthritis in the C3H mouse model.

## Results

### Overview of spatial transcriptomics workflow and quality controls

C3H mice were inoculated intradermally with 1×10^4^ *Bb*, which establishes an infection similar to natural tick transmission [9], or with media as a vehicle-treated control. The infection was resolved by treating mice intraperitoneally with 50mg/kg Doxycycline starting at 3 weeks post-infection. Joint tissue samples were harvested at two and four weeks post-infection, and subsequently processed for H&E and immunohistochemical (IHC) staining for Vimentin (VIM) and I-Eκ (Figs. 1 and 2), which are known markers of synoviocyte and *Bb* infection respectively [10].

**Figure 1:**
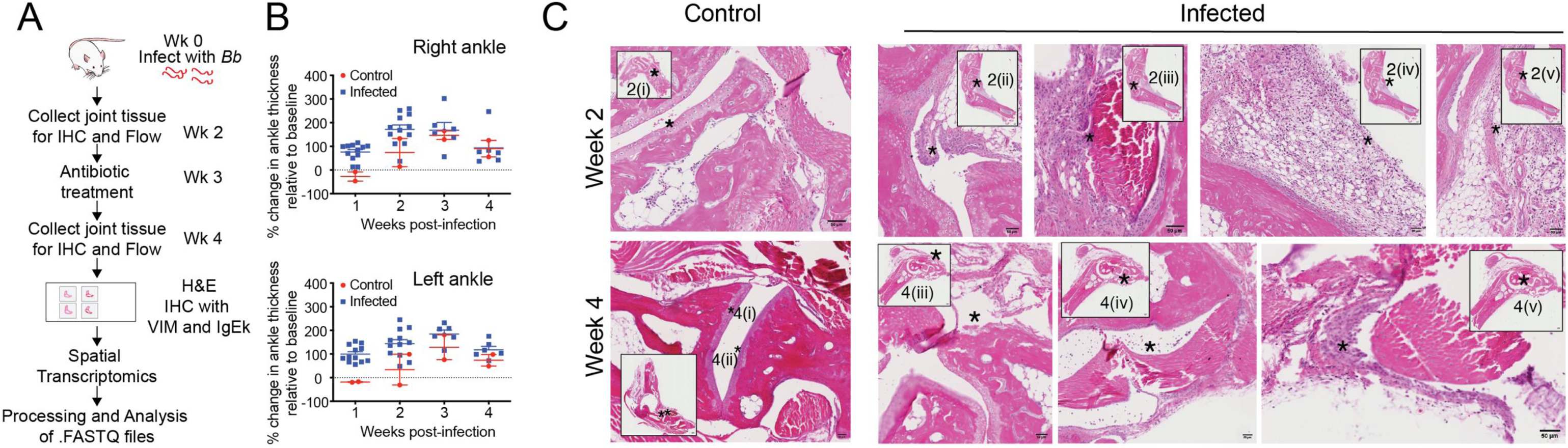
Overview of spatial transcriptomics workflow and quality controls. **(A)** Schematic of the experimental workflow. Mice were infected with *Borrelia burgdorferi* (*Bb*) at week 0. Antibiotic treatment was administered at week 3. Joint tissue was collected at weeks 2 and 4, sectioned and processed for hematoxylin and eosin (H&E) staining and immunohistochemistry (IHC) using antibodies against Vimentin (VIM) and IgEk. Tissue was also subjected to spatial transcriptomics, with downstream processing and analysis of FASTQ files. **(B)** Percent change in right ankle (top) and left ankle (bottom) thickness relative to baseline over 4 weeks post-infection in control (red circles) and *Bb*-infected (blue squares) mice. Data are shown as individual measurements with median and interquartile range. **(C)** Representative H&E-stained sections of ankle joints from uninfected (top rows) and *Bb*-infected (bottom rows) mice at week 2 (left columns) and week 4 (right columns). Arrows indicate areas of histopathological changes. Images are shown at equivalent magnification. 2(i)= normal articular cartilage with underlying bone, 2(ii)= papillary synovium, 2(iii)=synovial hyperplasia, 2(iv)= fibrin deposits and 2(v)=dense inflammation with lymphocytes, plasma. Week 4 images show 4(i and ii) = normal articular cartilage with underlying bone; 4(iii) =erosion of articular cartilage, 4(Iv)=mild inflammation, 4(v)= synovial hyperplasia.

**Figure 2.**
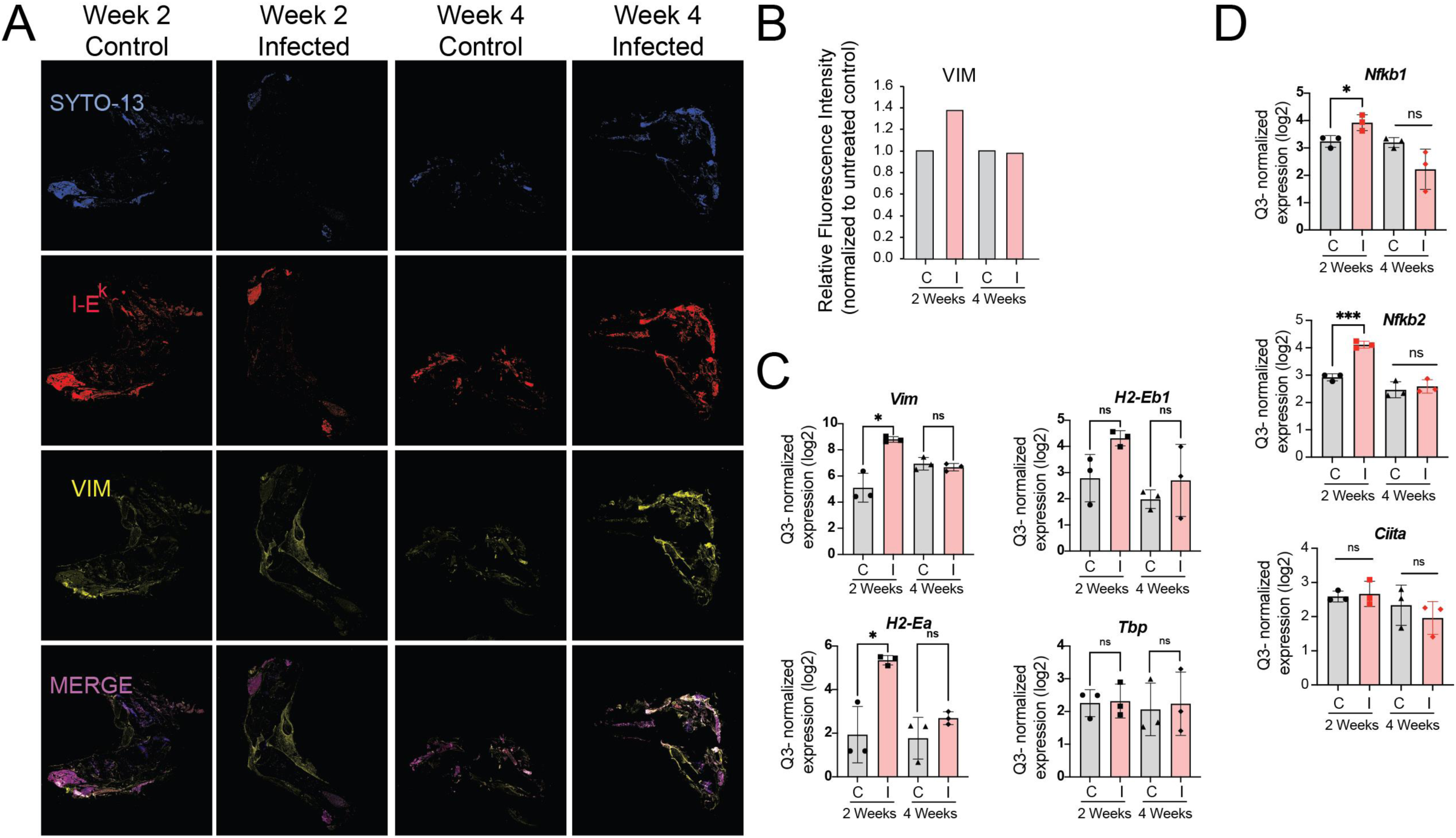
Immunofluorescence staining and gene expression of mesenchymal and immune markers in *Borrelia burgdorferi*-infected joint tissue. **(A)** Representative immunofluorescence images of ankle joint sections from uninfected and *Bb*-infected mice at weeks 2 and 4 post-infection. Tissue sections were stained with SYTO-13 (blue; nucleic acid counterstain), anti-I-Eᵏ (red; MHC class II marker), and anti-Vimentin (VIM; yellow). The bottom row shows a merged composite of all three channels. Columns represent the four experimental groups: Week 2 Uninfected, Week 2 Infected, Week 4 Uninfected, and Week 4 Infected. **(B)** Images were analyzed using ImageJ (NIH). Each channel was processed individually by first converting to 8-bit grayscale, followed by intensity thresholding to isolate signal from background. Bars represent normalized mean fluorescence intensity of VIM relative to nuclear stain (SYTO13). **(C)** Log₂ fold change expression of *Vim*, *H2-Ea*, *H2-Eb1*, and *Tbp* and **(D)** *Nfkb1*, *Nfkb2*, and *Ciita* in joint tissue from control (C, gray) and Bb-infected (I, pink) mice at 2 and 4 weeks post-infection. Individual data points represent biological replicates; bars represent mean ± standard deviation. Statistical comparisons between uninfected and infected groups at each timepoint were performed using an unpaired t-test; *p < 0.05, ns = not significant.

Resolution of infection following treatment with antibiotics was confirmed via ear-punch subcultures. Tibial tarsal joint thicknesses of both ankles were measured weekly starting one-week post-infection and plotted as percent change from baseline (Fig. 1B). Peak swelling occurred at one to two weeks post-infection, with around a 150% increase over baseline at 2 weeks, aligning with prior reports of maximal inflammation occurring around the 2-week time frame [1, 11]. By four weeks (one-week post-antibiotic treatment), swelling reduced to 100% of baseline and was statistically indistinguishable from controls (Fig. 1B).

Joints were harvested at two and four weeks post-infection, and H&E staining was performed (Fig. 1C). Compared to uninfected controls, infected mice at both time points showed clear evidence of joint inflammation. This included dense lymphocyte, plasma cell, and neutrophil infiltrates within the sub synovium (2(v)), synovial hyperplasia (2(iii), 4(v)), pronounced edema (2(v)), and focal fibrin deposition (2(v)). Foci of chronic papillary synovitis and articular cartilage destruction (4(iii)) were also observed.

Overall, joints collected two weeks post-infection were scored at a histological arthritis severity score of 4 while joints collected four weeks post infection received a score of 2. No markers of arthritis were annotated at two weeks or four weeks in control mice. Joints from both two- and four-week timepoints were selected for further detailed analysis to contrast peak inflammation with partial resolution. The two-week stage revealed acute synovitis driven by active infection, while the four-week stage (post-antibiotic) modeled persistent, immune-mediated arthritis. These data reflect the immunological mechanisms sustaining joint pathology in each stage of the infection.

### Immunofluorescence staining and spatial transcriptomic profiling of *Bb* -infected joint tissue

The joint tissues collected two and four weeks post infection were formalin-fixed, paraffin embedded, sectioned and stained with antibodies against I-E*^k^* (red) and VIM (yellow), while SYTO-13 (blue) was used to stain nuclei (Fig. 2A). High I-E*^k^* expression reflects upregulation of MHC class II on antigen-presenting cells and synovial stromal cells, supporting enhanced presentation of *Bb* antigens and sustained CD4⁺ T cell–mediated inflammation in LA [10]. VIM is an intermediate filament protein that becomes upregulated and redistributed in activated synoviocytes and infiltrating immune cells, where it contributes to synovial inflammation and tissue remodeling in pathological states such as rheumatoid arthritis [12, 13]. VIM was elevated in stained tissue from *Bb* infected mice relative to control mice at two weeks as expected (Fig. 2B).

Regions of synovium from the joint tissue of mice at two and four weeks post infection with high VIM and I-E*^k^* expression containing approximately 100 cells were then selected as regions of interest (ROIs), along with anatomically similar regions in the controls (Supplementary Fig. S1). Following UV-photocleavage of oligo-tagged probes and collection of regions-of-interest (ROIs) material, libraries were prepared according to the manufacturer’s protocol and sequenced and processed for downstream analysis.

Normalized probe intensities for *VIM* transcript levels (mean normalized counts of three ROIs) were significantly elevated at two weeks post infection compared with both control and the four-week post-infection groups (Fig. 2C), consistent with the increase in VIM protein expression observed in the immunohistochemically stained joint tissue. The MHC class II genes *H2-Ea* and *H2-Eb1*, encoding the α and β chains of the I-E molecule, were also examined. Normalized expression (log2) values showed that *H2-Ea* was significantly increased two weeks after infection relative to uninfected controls, while no significant increase was noted at four weeks post infection (Fig. 2C). On the other hand, *H2-Eb1* did not differ significantly relative to uninfected controls at both two and four weeks post-infection, suggesting selective regulation of *I-E* chain-related gene expression at peak inflammation (Fig. 2C). While MHC class II upregulation is documented in C3H Lyme arthritis model [14, 15], there are currently no studies reporting non-selective increases in both *H2-Ea* and *H2-Eb1* in Lyme arthritis. As a control, we surveyed *Tbp* (TATA-box binding protein) probe intensities (Fig. 2C) because it has been identified as one of the most stable reference genes in collagen-induced and spontaneous arthritis in mouse joints [16]. As expected, no significant differences were found between control and infected joints. As the alpha and beta chains of the *I-E* allele are coordinately regulated [17], it is possible that the selective significant upregulation of *H2-Ea* could be likely due to other transcription factors independent of CIIT-a, for instance, the NF-kB signaling pathway members [18]. Indeed, probes for both *Nfkb1* and *Nfkb2* are significantly upregulated in *Bb*-infected relative to uninfected tissue at 2 weeks, but not at four weeks post infection; while *Ciita* levels are not significantly different (Fig. 2D).

### Temporally distinct host gene expression and pathway enrichment profiles during *Bb* infection

To further resolve temporal transcriptional differences, three regions representing three anatomically similar areas of synovium from control and infected mice two and four weeks after inoculation were compared. At two weeks post-infection, infected and uninfected joints exhibited distinct expression profiles, with a clear separation between upregulated and downregulated genes (Fig. 3A). By four weeks post infection, these differences were less pronounced, with no identical genes appearing in both heatmaps, suggesting the host gene expression profile is dependent on infection stage.

**Figure 3:**
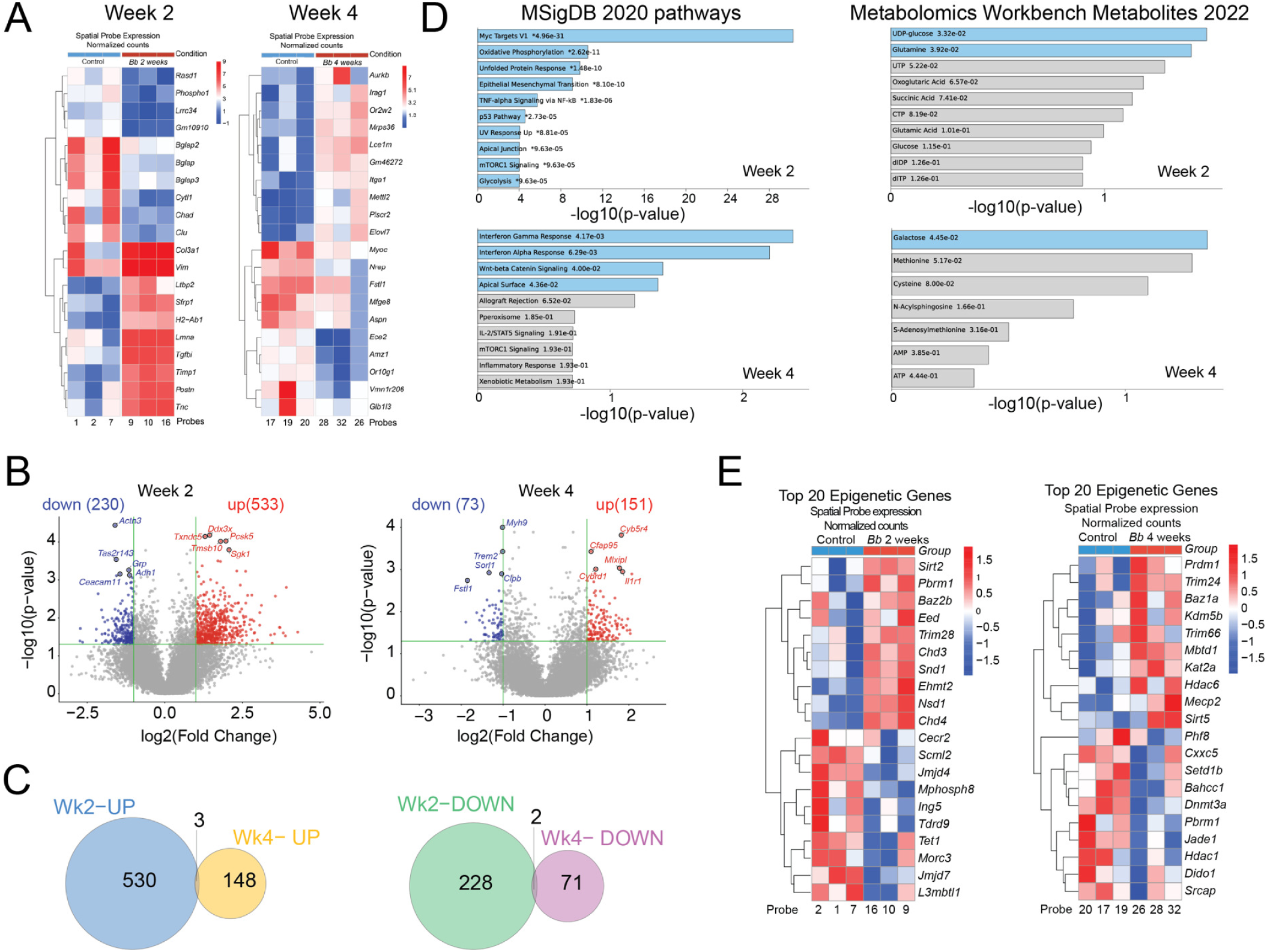
Spatial transcriptomics data analysis reveals changes in immune regulatory genes, signaling pathways, and extracellular matrix proteins. (A) Heatmap displaying log₂ Q3-normalized counts for the top 10 upregulated and top 10 downregulated genes in *Borrelia burgdorferi*-infected (*Bb* 2 weeks; probes 9, 10, 16; Bb 4 weeks; probes 28,32,26) versus vehicle-treated (2 weeks; probes 1, 2, 7 and 4 weeks; probes 17,10,20) and tissue sections, as measured by GeoMx Digital Spatial Profiling (DSP). Genes were ranked by mean expression difference (infected − vehicle) and the top 10 genes in each direction were selected for display. Rows are hierarchically clustered (Euclidean distance, complete linkage); columns are ordered by condition. Color scale represents normalized expression values from blue (low) to red (high). (B) Volcano plots of significant up (red) or down (blue)-regulated genes at week 2 (left) and week 4 (right) after *Bb* infection relative to vehicle-treated samples; (C) Number of overlapping genes between weeks 2 and 4 that are upregulated (top Venn diagram) and downregulated (bottom Venn); (D) Top enriched MSigDB Hallmark pathways (left) and Metabolomics metabolites (right) in the upregulated genes at week 2 (top two graphs) and week 4 (bottom two graphs) after *Bb* infection. X axis indicates –log10(p-value) respectively were determined using EnrichR and Appyter. (E) Expression profiles of the 10 most upregulated and 10 most downregulated epigenetic genes between Control two weeks (probes 1,2,7) and two week *Bb* infected (probes 9,10,16, left panel); and control four weeks (probes 17,19, and 20) and *Bb* infected at four weeks (probes 26, 28, and 32), right panel. Expression values are row-scaled (z-score), with hierarchical clustering applied to both genes and samples.

Differentially expressed genes at 2 and 4 weeks were visualized using volcano plots, with a p-value cutoff of 0.05 and an absolute log_2_ fold-change threshold of 2 (Fig. 3B). At two weeks post infection, 533 genes were significantly upregulated and 230 were significantly downregulated. At four weeks post infection, these numbers were reduced, with 151 significantly upregulated and 73 significantly downregulated genes. Few overlaps between the two- and four- week time points were observed, with only 2 overlapping downregulated genes (*Or5b123*, *Spaca7b*) and 3 overlapping upregulated genes (*Abcc5*, *Pde4b*, *Snx20)* (Fig. 3C). This comparison highlights temporal reprogramming of the joint transcriptome between peak and resolving LA.

Next, we performed Gene Ontology (GO) analysis to identify the most significantly enriched MSigDB Hallmark 2020 pathways and metabolomic metabolites, utilizing the EnrichR analytical framework [19] (Fig. 3D). At two weeks post-infection, many top pathways aligned with known Lyme arthritis mechanisms such as oxidative phosphorylation, epithelial-mesenchymal transition (EMT), glycolysis and NF-κB signaling were upregulated. Synovial fibroblasts and macrophages are known to undergo glycolysis and oxidative phosphorylation reprogramming in similar arthritic pathologies like rheumatoid arthritis, driven by NF-κB and TNFα signaling [20], further confirming that the selected regions capture LA infection relevant biology. Downregulated pathways at week two included Myc targets, EMT, oxidative phosphorylation, and PI3K/AKT/mTOR signaling (Supplementary Fig. S2A). While a direct connection between MYC and LA is not clear, MYC is known to be elevated in arthritis [21] and impacts metabolic programming of regulatory T-cells [22], including oxidative phosphorylation.

At four weeks post-infection (one week after antibiotic treatment) MSigdB GO pathway analysis shows enrichment in IFNγ response, Wnt-β-catenin signaling, IL-2/STAT signaling, apical junction signaling and inflammatory response pathways, indicating a persistent immune activation and tissue repair phase (Fig. 3D). While IFNγ and inflammatory responses reflect ongoing T-cell and macrophage activity against residual *Bb* antigens [23], Wnt-β-catenin signaling suggested epithelial progenitor proliferation and EMT post-damage driving epithelial and stromal regeneration [24]. Interferon signaling and inflammatory response pathways were downregulated at four weeks (Supplementary Fig. S2B), consistent with the resolution of the infection by this time.

Additionally, when metabolic pathways were surveyed via Enrichr, distinct metabolic pathways were upregulated between two and four weeks post-infection (Fig. 3D), with glucose metabolism upregulated at two weeks and galactose pathways at week four. Although not significant, there is a slight increase in methionine and cysteine pathways at week four (Fig. 3D). Downregulated pathways at week two included genes involved in UTP and CTP metabolism, and galactose and methionine metabolism by week four (Supplementary Fig. 3D). This temporal shift likely reflects an early glutamine-driven metabolism, supporting quick energy and biosynthesis in proliferating innate immune cells [25], while later cysteine and methionine pathways bolster adaptive T-cell responses and mitigate ROS [26] at four weeks post infection.

Metabolic reprogramming during Lyme disease is not unique to synovial tissue, as multiple studies demonstrate broad host metabolic shifts. *Bb* infection strongly upregulates glutathione pathways in human macrophages and PBMCs, which are essential for cytokine production via glutathionylation [27]. Similarly, profiling the serum metabolome in early Lyme disease reveals enrichment in cysteine and glutathione metabolism [28]. These metabolic adaptations parallel the glutamine and cysteine enriched shifts observed in our GO analysis, supporting a unified model of temporal immune-metabolic progression in LA.

Since metabolites involved in methionine pathways and acetyl-CoA link metabolism to epigenetic regulation, we investigated whether genes coding for epigenetic machinery were differentially expressed between control and *Bb* infected tissues (E). Strikingly, genes involved in transcriptional silencing were upregulated at two weeks post-infection, including *Ehmt2, Chd3, Chd4, Baz2b, Eed*, and *Sirt2* (Fig. 3E, left panel). Notably, *Chd3* and *Chd4* are core ATPase components of the Mi-2/NuRD chromatin remodeling complex, suggesting coordinated repression of immune gene programs early during infection. At four weeks post-infection, epigenetic regulators including *Baz1a*, *Mecp2, Sirt5, Kdm5b, Kat2a, Hdac6*, and *Prdm1* (Fig. 3E, right panel) were upregulated in *Bb*-infected tissues relative to control. The upregulation of *Prdm1* (*Blimp-1*), a key regulator of T-cell exhaustion in chronic infections [29], suggests that the adaptive immune response may be entering an exhausted or tolerized state by week four.

### Distinct temporal Immune pathway-related host responses to *Bb*

To probe differences between two and four weeks post-infection, we used pathway analysis datasets generated by Reactome (Build 78 + NCBI_08122021) within the GeoMx DSP Analysis Suite. The top 10 immune pathways were ranked by normalized enrichment score using Reactome for both two and four weeks post- infection (Fig. 4A) and visualized as a bubble plot. Pathway analysis revealed programs such as B-cell receptor signaling, NF-κB activation in B- cells, and antigen cross-presentation as dominant at two weeks post-infection, reflecting acute, antigen-driven synovitis with robust humoral responses [10]. By four weeks post-antibiotic treatment, pathways shifted to CD28 co-stimulation, CD28-dependent PI3K/Akt signaling, and IFN-stimulated gene activation, however, these pathways were below the threshold for significance. This temporal progression highlights a transition from infection-centric inflammation to a persistent, immune-mediated arthritic state akin to post-treatment LA.

**Figure 4:**
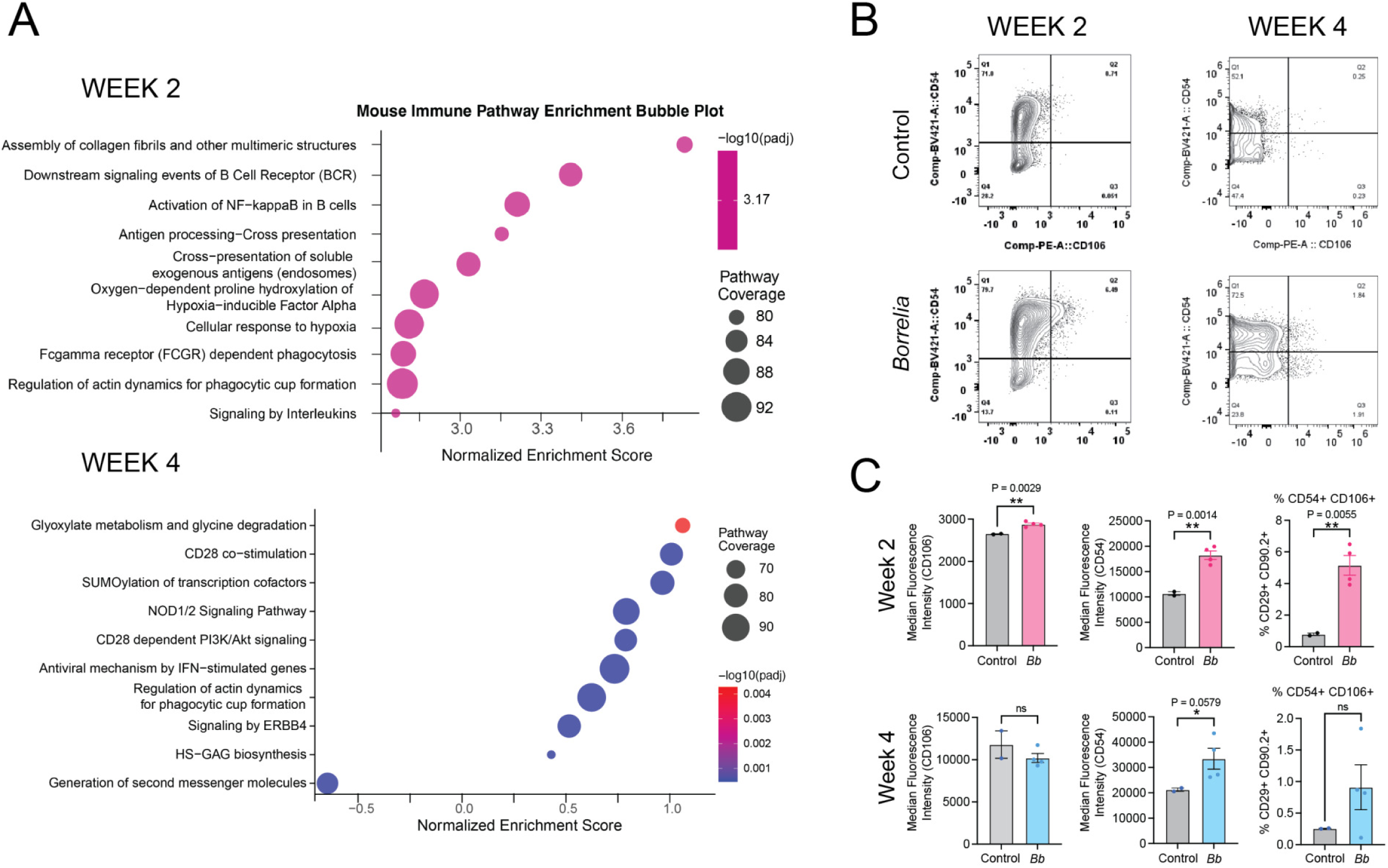
Immune pathway responses to *Borrelia burgdorferi*. (A) (B) (C) Increased frequency of VCAM-1/ICAM-1 cells in the synovial stromal compartment at week 2 post infection. Bar graph showing the frequency of CD106 (VCAM-1) and CD54 (ICAM-1) double positive cells expressed as % of CD29 (Integrin-b1) and CD90.2 (Thy1.2) double positive cells. Error bars represent SEM, n=2-4. Individual mouse values are shown as scatter points. *p<0.05, Welch’s Unpaired t-test was performed using GraphPad Prism version 11.0.0.

Next, we asked if synovial markers were altered in a temporal fashion in mice infected with *Bb*. To do this, we used mouse joint samples isolated at weeks two and four in parallel with the tissue isolation for spatial transcriptomics to compare changes in CD54^+^ and CD106^+^ populations over time in control versus *Bb*-infected samples (Fig. 4B) using flow cytometry. CD54, also known as Intracellular Adhesion Molecule 1 (ICAM-1), is a key mediator of leukocyte adhesion and activation during inflammatory responses [30]. CD106 (vascular cell adhesion molecule-1; VCAM-1) marks a subset of mesenchymal and endothelial cells and facilitates firm adhesion of immune cells to the endothelium [31]. In the synovia of LA patients, ICAM-1 is strongly upregulated on lining and endothelial cells, while VCAM-1 shows moderate upregulating supporting sustained leukocyte trafficking into the synovium [32].

At week two post-infection, the joints of both VT and *Bb*-infected C3H mice were dominated by CD54^+^ and CD106^-^ cells; however, *Bb*-treated samples exhibited a modest increase in CD106⁺ populations relative to controls (Fig. 4B). By week four, *Bb*-infected samples show a clear expansion of CD54^+^ and CD106^+^ double-positive population compared to the controls. To quantify these changes, we next assessed median fluorescence intensity (MFI) of CD54 and CD106, as well as the frequency of CD54⁺/CD106⁺ double-positive cells (Fig. 4C). At week two, *Bb*-infected samples showed a significant increase in both CD106 and CD54 expression relative to controls (Fig. 4C top right panel). Consistent with this, the proportion of CD54⁺/CD106⁺ cells was also significantly elevated, confirming an early induction of an activated, adhesion molecule - high phenotype. By week four, CD106 expression was no longer significantly different between groups (Fig. 4C bottom left panel), while CD54 expression showed a continued upward trend in *Bb*-treated samples (Fig. 4C bottom middle panel). Although the frequency of CD54⁺CD106⁺ cells remained higher in *Bb*-treated samples at this time point, this difference did not reach statistical significance.

Together, these data indicate that *Bb* infection induces an early and robust upregulation of both ICAM-1 and VCAM-1, which partially persists but becomes more variable over time. This temporal pattern suggests that peak endothelial and synovial activation occurs during the early phase of infection, potentially driving the activation of NF-κB and downstream interleukin signaling seen in the pathway analysis (Fig. 4A). ICAM-1 and VCAM-1 promoters both contain NF-κB response elements and interleukin signaling (especially IL-1, IL-6 and IL-17) strongly induces endothelial activation [33–35]. Finally, a more heterogeneous or regulated adhesion molecule expression can be seen at later stages.

### Comparison of medial and lateral humeroulnar synovium reveal distinct spatially localized host responses to *Bb* infection

Regional differences in transcriptomics have been annotated in arthritic synovial tissue. For instance, fibroblasts in the lining of rheumatoid arthritic lesions were found to express more genes related to inflammation such as degradation of collagen and extracellular matrix proteins and Interleukin signaling, while cells in the sub-lining expressed genes related to adaptive immunity such as antigen processing and presentation [36]. A study investigating juvenile idiopathic arthritis also found regional differences in the spatial transcriptomic joint profile, identifying microanatomical structures with their own unique transcriptional signatures [37]. Although no studies to date have compared regional differences in gene expression in Lyme arthritis, different regional fibroblast and macrophage gradients near varying lymphoid aggregates have been noted as Lyme synovitis forms in patients [38].

To explore regional differences in the spatial transcriptomic profiles during peak inflammation within the same tissue sample, we compared the synovium located near the medial humeroulnar joint to synovium located near the lateral humeroulnar joint (Fig. 5A). We noted 514 upregulated genes in the medial and 207 in the lateral synovium (log_2_fc >2, p-value <0.05) with only 19 overlapping genes. Additionally, there are 225 significantly downregulated genes in the medial and 92 in the lateral synovium with 5 overlaps (Fig. 5B), highlighting spatially compartmentalized host responses to *Bb* infection. The top 10 up- and top 10 down-regulated genes from probes in the lateral and medial synovium were plotted in a heatmap (Fig. 5C).

**Figure 5.**
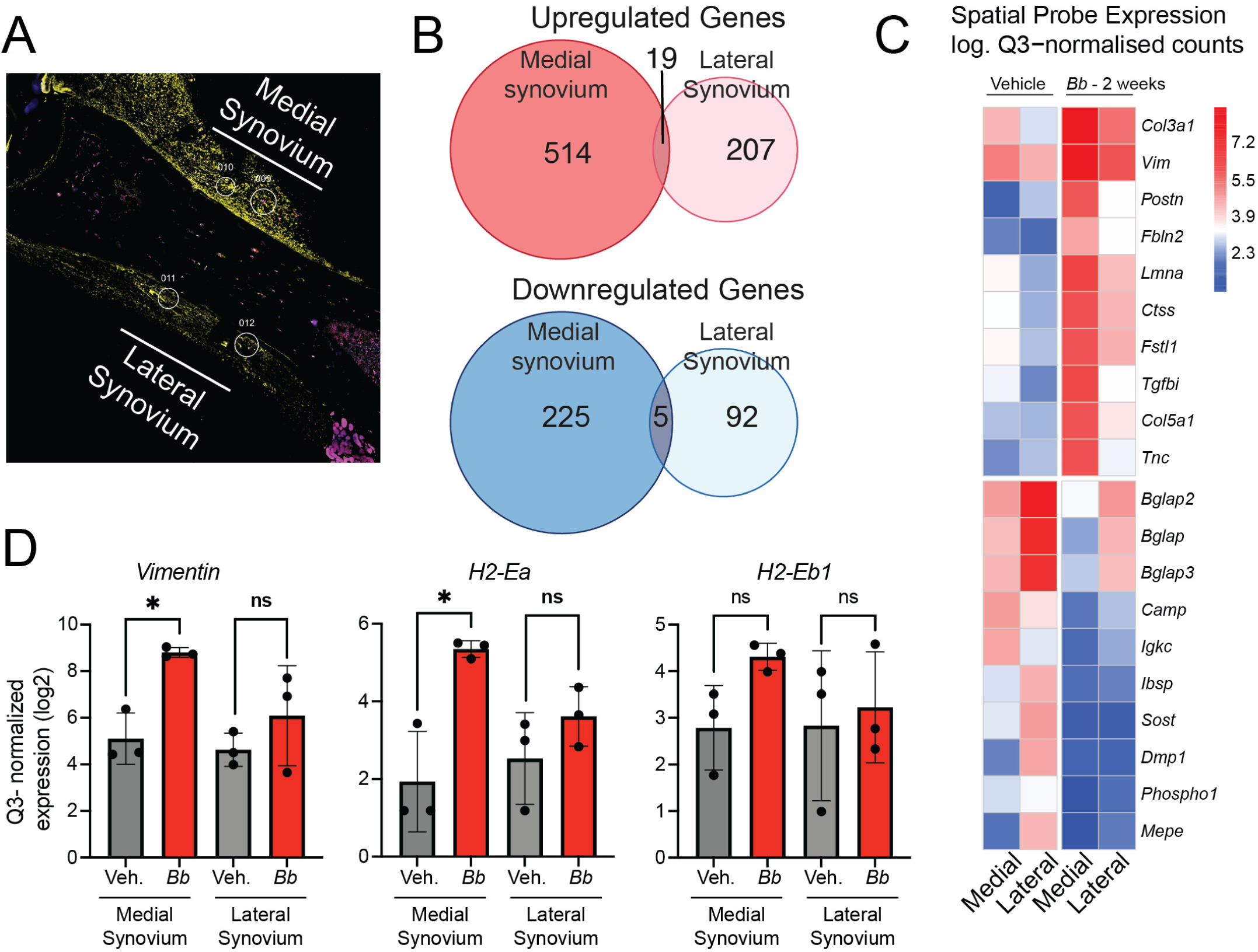
Comparison of synoviocyte-rich and Synoviocyte-bone border regions during *Borrelia* infection. (A) (B) (C)Heatmap displaying the top 10 up- and top 10 down-regulated genes comparing probes in the synovium region with probes in the bone border region at two weeks post-infection with *Bb*. Expression values represent log₂-transformed, Q3-normalised counts averaged across biological replicates within each group. Genes were ranked by the average log₂ fold change (infected − uninfected) across both regions, and the top 10 most upregulated and top 10 most downregulated genes are shown. Gene names are displayed on the y-axis; color scale reflects expression level from low (blue) to high (red). (D) Q3 normalized expression average of 3 probes from lateral and medial synovium-rich regions plotted for *Vim*, *H2-Ea*, *H2-Eb1* genes. *= p-value<0.05, ns= not significant.

Although the same groups of genes were up/down regulated in the infected relative to the control, these differences were amplified in the medial regions of the joint relative to the lateral regions. Additionally, *VIM* transcript levels were significantly elevated in the medial humeroulnar regions of synovium relative to lateral regions of synovium (Fig. 5D, first panel). Similarly, the MHC class II genes *H2-Ea* was significantly increased in the interior region 2 weeks after infection relative to all other groups (Fig. 5D, middle panel), while *H2-Eb1* was not significantly elevated in any group (Fig. 5D, last panel). Overall, the medial humeroulnar region appears to have a more polarized and synchronized gene expression relative to the lateral region.

### Concordance between spatial transcriptomics and scRNA-seq synoviocyte gene expression changes following *Bb* infection

Next, we compared our spatial transcriptomics data with published scRNA-seq [39] in order to compare the broad transcriptomic depth of scRNA-seq with the tissue context and positional information of our spatial datasets. The scRNA-seq data were obtained from joints of mice infected with *Bb* and collected at weeks two and four, similar to our data. We reasoned that the integration of synoviocyte clusters in the scRNAseq dataset with our spatial dataset would enable us to identify distinct cell populations and their gene expression profiles, and also would determine where those cells reside within the tissue architecture of the joint. To do so, we compared gene expression profiles in our spatial data with synoviocyte cluster from the published scRNAseq dataset.

A major caveat to note here is that different mouse strains were used in generating these datasets: Spatial transcriptomic profiling was performed in C3H mice, while scRNAseq data were derived from synoviocytes isolated from C57BL/6 mice. Given this cross-strain experimental design, we reasoned that genes exhibiting concordant directional changes across both modalities may be the most biologically robust, as they represent responses conserved across distinct genetic backgrounds and measurement technologies. A second caveat associated with this comparison is that neither dataset had reliable p-values (spatial transcriptomics data has low n, so FDR is challenging; while scRNAseq p-values are inflated by pseudo replication due to n=1). However, we felt that comparing purely based on log2 fold changes could still be a valuable way to explore the two datasets and potentially generate new hypotheses. With these caveats in mind, we assessed the degree of concordance between spatial transcriptomics and single-cell RNA sequencing (scRNA-seq) data from synoviocytes following *Bb* infection for two and four weeks (Fig. 6A). This integrative approach enabled us to determine whether transcriptional changes observed at the tissue level are recapitulated within individual clusters of synoviocyte cells.

**Figure 6.**
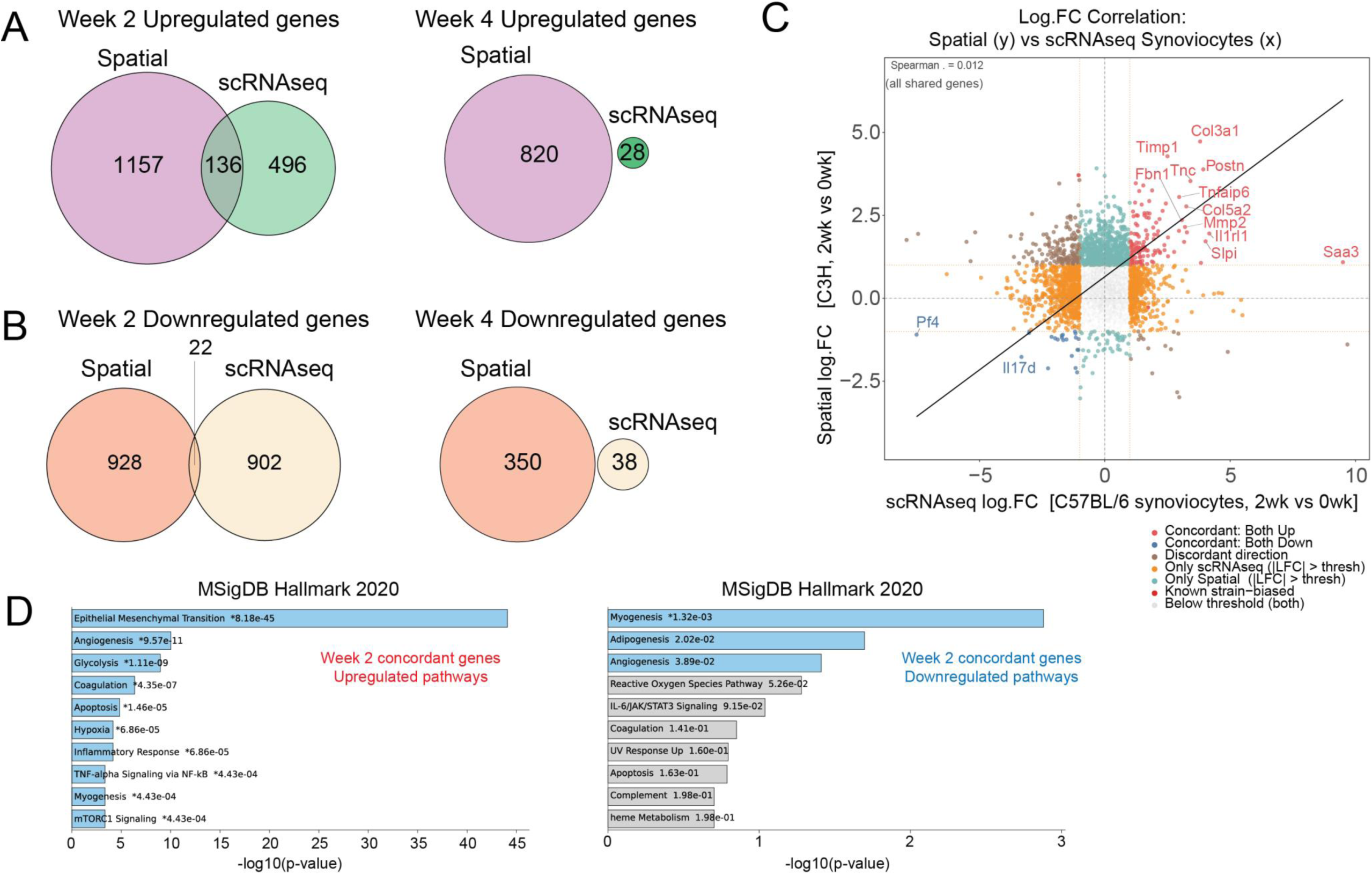
Concordance between spatial transcriptomics and scRNA-seq synoviocyte gene expression changes following *Bb* infection. (**A**) Venn diagrams depicting the overlap of upregulated genes identified by spatial transcriptomics and scRNA-seq synoviocyte cluster data at Week 2 (left) and Week 4 (right) post-*Bb* infection. Numbers indicate genes unique to each modality or shared between both. (**B**) Venn diagrams showing the overlap of downregulated genes from the same comparisons at Week 2 (left) and Week 4 (right). (**C**) Scatter plot of log fold-change (log.FC) correlation between spatial transcriptomics (y-axis; C3H strain, 2 weeks vs. 0 weeks) and scRNA-seq synoviocytes (x-axis; C57BL/6 strain, 2 weeks vs. 0 weeks) for all shared genes. Each point represents a gene, colored by concordance category: concordant upregulation in both modalities (red), concordant downregulation in both modalities (blue), discordant direction between modalities (brown), significant only in scRNA-seq (|log.FC| > threshold; orange), significant only in spatial transcriptomics (|log.FC| > threshold; teal), known strain-biased genes (pink), or below threshold in both modalities (gray). Select genes of interest are labeled. The diagonal line represents the line of best fit. (**D**) Bar plots showing MSigDB pathway enrichment (via EnrichR/Appyter) for Week 2 concordant upregulated genes (left) and concordant downregulated genes (right), defined as genes significantly differentially expressed in the same direction in both spatial transcriptomics and scRNA-seq synoviocyte data. Pathways are ranked by −log₁₀(p-value); asterisks (*) denote pathways meeting a significance threshold. Blue bars indicate pathways passing the significance cutoff; gray bars indicate nominally enriched pathways.

At week two post-infection, we observed an overlap of 136 concordantly upregulated genes between spatial transcriptomics and scRNA-seq datasets (Fig. 6A). However, a large proportion of genes remained distinctly different between the scRNA-seq and spatial datasets, possibly reflecting the broader cellular net cast by spatial transcriptomics relative to the more restricted resolution of scRNA-seq. By week four, concordance in upregulated genes was reduced to zero using our threshold of |log₂FC| > 1.0 (fold-change > 2). A similar pattern was observed for downregulated genes (Fig. 6B), where the overlap was modest at week two with 22 concordantly downregulated genes observed.

By week four, however, there were no overlapping genes that were concordantly downregulated with these parameters. As the scRNAseq synoviocyte cluster data at week four only had 1,033 genes altered, vs. 5,274 in week two, this might potentially explain in part why the overlap was nonexistent.

We visualized the magnitude and consistency of fold-changes across both modalities at the same time in a different way by generating a dot plot for the top concordant genes at week two relative to uninfected controls, ranked by mean absolute log₂FC across both datasets (Fig. 6C). This again revealed a weak but positive correlation between spatial and scRNA-seq expression changes, indicating partial agreement in the direction and magnitude of transcriptional responses. Notably, a subset of genes demonstrated concordant upregulation across both platforms, including several associated with extracellular matrix remodeling (*Col1a1, Col3a1, Mmp2, Timp1*), fibroblast activation (*Postn, Postn, Runx1, Ctnnb1*), and inflammation (*Saa3, Ccl7, Tnfaip6*), while others exhibited discordant behavior, potentially reflecting strain-specific effects (C3H vs. C57BL/6) or differences in cellular composition between datasets.

Finally, pathway enrichment analysis of concordantly regulated genes highlighted biologically relevant processes associated with infection. Concordant upregulated genes at week two were enriched for pathways related to cytoskeletal organization, integrin signaling, and extracellular matrix receptor interactions, consistent with activation of stromal and adhesion programs (Fig. 6D, left panel). In contrast, concordant downregulated genes were associated with metabolic and cytokine signaling pathways, suggesting a coordinated shift in synoviocyte functional states during early infection (Fig. 6D, right panel). Together, these findings demonstrate that *Bb* infection induces partially conserved transcriptional programs between spatial joint profiling and synoviocyte-specific scRNA-seq, with the strongest concordance observed during early infection.

## Discussion

Spatial transcriptomics provides unique benefits for investigating subacute and post-treatment Lyme arthritis, as it maintains tissue structure while detecting spatially restricted immune activity overlooked by bulk and single-cell RNA-seq. Although previous transcriptomic studies of Lyme arthritis have highlighted disrupted immune circuits [40–42], they depended on bulk sequencing that blends signals from heterogeneous cell populations and microenvironments, masking localized gene expression dynamics. By contrast, spatial transcriptomics delivers comprehensive, architecture-preserving gene expression maps [6], illuminating cellular neighborhoods and transcriptional hotspots that drive synovial inflammation. In this study, we combined histopathology, spatial transcriptomics, and flow cytometry to define the temporal and spatial dynamics of joint inflammation during *Bb* infection. Our findings reveal a coordinated progression from acute, infection-driven synovitis at two weeks to a more heterogeneous, immune-mediated state following antibiotic treatment at four weeks, providing insight into mechanisms that may guide persistent inflammation in LA.

Consistent with prior reports, we observed peak joint swelling and histopathological severity at two weeks post-infection (Fig. 1B and C), characterized by robust immune cell infiltration, synoviocyte hyperplasia, and tissue remodeling [11, 32]. We demonstrate that VIM and I-E^k^ immunostaining correspond directly with increased synovial transcript levels of *Vim* and *I-E^k^* related gene expression, further validating the biology of LA in this model (Fig. 2). Additionally, the selective upregulation of *H2-Ea* but not *H2-Eb1* (Fig. 2B) suggests nuanced regulation of MHC class II components during Lyme arthritis, which may influence antigen presentation and T-cell activation in a gene-specific manner, potentially due to NFKB family members (Fig. 2C).

Spatial transcriptomics analysis mirrored the histopathological findings. Our analysis revealed stage-specific transcriptional reprogramming, with 533 upregulated and 230 downregulated genes during acute inflammation at week two, compared to 151 upregulated and 73 downregulated genes during the resolution phase at week four (Fig. 3). The data also revealed stage-specific transcriptional reprogramming, with immune pathway enrichment dominated by B-cell signaling, NF-κB activation, and antigen cross-presentation, consistent with acute humoral immune responses (Fig. 4). In contrast, the week four time point was characterized by CD28 co-stimulation, PI3K/Akt signaling, and interferon-stimulated gene expression, reflecting persistent T-cell activation and chronic inflammatory signaling (Fig. 4). Although antibiotic treatment reduced overt inflammation by four weeks, residual histological and transcriptional changes persisted (Fig. 3B), supporting the concept that Lyme arthritis can transition from an infection driven pathology to a more immune- mediated pathology. This is further reinforced by our pathway analyses, which show a shift from early innate and humoral immune activation including NF-κB signaling, B-cell receptor signaling, and metabolic reprogramming to later-stage pathways dominated by T-cell co-stimulation, interferon signaling, and tissue repair processes (Fig. 4A).

At the cellular level, flow cytometry analysis (Fig. 3B, C) revealed early upregulation of adhesion molecules ICAM-1 (CD54) and VCAM-1 (CD106), consistent with enhanced leukocyte recruitment during acute infection [43], which is consistent with the H&E staining showing increased leucocyte infiltration (Fig. 2). While this adhesion molecule–high phenotype was most pronounced at two weeks, partial persistence at four weeks suggests continued, albeit variable, endothelial and stromal activation. These findings parallel observations in human Lyme arthritis synovium [32] and provide functional evidence linking transcriptional changes to cellular phenotypes that facilitate immune cell trafficking.

Our data also demonstrate that *Bb* infection induces significant metabolic reprogramming within the joint. Early enrichment of glycolytic and oxidative phosphorylation pathways, alongside glutamine-associated metabolism (Fig. 3D), is consistent with the energetic demands of rapidly proliferating and activated immune cells [44]. In contrast, later enrichment of cysteine and glutathione-related pathways suggests a shift toward redox regulation and resolution-phase metabolism (Fig. 3D). These findings align with prior studies in peripheral immune cells [27, 28] and extend them to the joint microenvironment, highlighting metabolism as a key regulator of Lyme arthritis progression.

In addition to metabolic programming, we found several key epigenetic proteins involved in gene silencing were upregulated in *Bb*-infected tissues (Fig. 3E). While a previous study has implicated the Mi-2/NuRD complex in anti-microbial defense in *Drosophila* [45], to our knowledge, this is the first report of Mi-2/NuRD complex involvement in mammalian host responses to *Bb* infection. While *Ehmt2* upregulation has been documented in Borrelia-infected ticks [45, 46], the coordinated upregulation of the NuRD complex components (high levels of *Chd3/Chd4* at week 2, and high levels of *Mecp2* at week 4), specifically in infected mammalian tissues has not previously been reported. Interestingly, the Mi2-Nurd complex proteins are coordinately upregulated with *Prdm1*, contrary to previous reports that the Mi2-NuRD complex inhibits *Prdm1* expression in germinal center B-cells [47], suggesting tissue context and/or infection might be relevant for their interactions. Upregulation of the Mi2/NuRD complex proteins is mechanistically significant for understanding how *Bb* may manipulate host chromatin to establish tolerance and/or persistent responses to infection.

We also identify pronounced intra-joint heterogeneity, with the medial humeroulnar synovium exhibiting elevated *Vim* and *H2-Ea* expression (Figs. 5A and D), indicative of a regional specialization that mirrors observations in other inflammatory arthritis pathologies, where anatomical niches within the joint exhibit distinct functional programs [36, 37]. The medial synovium also demonstrated a more polarized, pronounced, and coordinated transcriptional response compared to the lateral regions, with minimal overlap in differentially expressed genes (Fig. 5B). These findings support an emerging model in which synoviocytes are not merely structural cells but active participants in shaping local immune responses. Joint cells such as myeloid cells, endothelial cells and fibroblasts are all able to propagate the robust IFN response in mouse models of Lyme arthritis [48]. Similarly, our results also suggest that localized microenvironments within the joint may differentially contribute to disease severity and persistence in Lyme arthritis. Supporting this idea, Lochhead et al. demonstrated through lineage sorting of Lyme-infected joint tissue that macrophage-like synoviocytes initiate type I IFN production in response to *Bb*, while CD45⁻CD31⁺ endothelial cells and fibroblasts amplify this response and serve as the primary sources of chemokines driving inflammatory cell recruitment [48]. Their data and ours highlight how functionally distinct cell populations within the joint microenvironment differentially shape disease.

Integration of spatial transcriptomics with scRNA-seq synoviocyte data [30] further demonstrated that *Bb* infection induces partially conserved transcriptional programs across platforms, particularly during early infection. The modest overlap in gene expression (Fig. 6A-B) and the positive correlation in fold-change values at two weeks (Fig. 6C) indicate that key inflammatory and extracellular matrix remodeling pathways are shared between tissue-level and cell-specific analyses. However, the marked reduction in concordance by four weeks (Fig. 6C) underscores increasing cellular heterogeneity and context-dependent regulation as inflammation evolves. Differences in mouse strain and cellular composition likely also contribute to this divergence, as the scRNA-seq data are from C57BL/6 mice, which are more resistant to *Borrelia* [49]. These findings demonstrate that spatial transcriptomics, especially when integrated with single-cell approaches, provides unique insight into the temporal and regional complexity of Lyme arthritis in the C3H mouse model.

Together, these findings support a model in which early Lyme arthritis is driven by coordinated activation of synoviocytes, endothelial cells, and infiltrating immune populations, resulting in robust inflammatory and metabolic reprogramming. As infection resolves, the joint transitions to a more heterogeneous state characterized by persistent immune signaling, synovial remodeling, and spatially distinct microenvironments that may sustain pathology independent of active infection. Collectively, our data support the idea of Lyme arthritis as a spatially and temporally evolving disease, in which host-driven inflammation affects joint pathology long after bacterial clearance. These observations lay the groundwork for identifying spatially resolved therapeutic targets capable of disrupting persistent synovial pathology in the post-infectious disease state.

## Materials and methods

### Bacterial strains and culture conditions

*B. burgdorferi* isolate B31-A3 was provided as a gift from Brian Stevenson, via Patricia Rosa [50]. Prior to all experiments, spirochetes were cultured to mid-log phase at 37°C under 5% CO2 in BSK-II culture medium supplemented with 6% rabbit serum and quantified by dark field microscopy using a Petroff-Hausser chamber.

### Infection and recovery of *Borrelia* from mice

All animal infections were carried out in accordance with approved protocols from the Institutional Animal Care and Use Committee (IACUC) of the University of North Dakota. C3HeB mice were purchased from The Jackson Laboratory (stock no. 000658). For all experiments, 6–8-week-old female mice were used and housed in groups of four. For infections, animals were placed under anesthesia using isoflurane inhalation, followed by subdermal inoculation in the dorsal thoracic midline with 100 uL of BSK-II medium containing 10^4^*Bb*. Two and four weeks following challenge by needle inoculation, mice were assayed for infectivity by culturing ear biopsy specimens for *B. burgdorferi* in BSK-II media [51]. Ear biopsy cultures were observed microscopically after a 7-day incubation for the presence of spirochetes and negative culture samples were maintained and observed for about 4 weeks before being discarded. Mice were euthanized at both 2 and 4 weeks post-infection for collection of the tibiotarsal joint, which was immediately snap frozen in liquid nitrogen for later imaging. Measurements of the tibiotarsal joint were taken every week over the course of 4 weeks.

### Tissue joint preparation

Tissue samples were fixed in formalin in 15 mL falcon tubes for 2-3 days, with the formalin solution replaced once on the second day. Following fixation, samples were washed twice with 12 mL PBS and resuspended in 12 mL of 20% EDTA (pH 7.4) for decalcification. Samples were placed on a shaker at 4°C, and the EDTA solution was refreshed every 2-3 days for two weeks, using a 20-fold excess volume to ensure complete saturation. After two weeks of decalcification, samples were washed three times with PBS, transferred to 70% ethanol (prepared with MilliQ water) for storage, and subsequently submitted for paraffin embedding at the University of North Dakota’s histology core.

Bone sections were cut at 4 μm thickness to optimize adhesion to slides. Leica Bond Plus slides were used as per the manufacturer’s instructions to mount the tissues. Tissues were maintained flat and kept on ice prior to sectioning, and cut on a cold block to ensure flat, transparent sections. Following sectioning, slides were placed on a heat block at 37-40°C overnight to partially melt the paraffin. The next morning, slides were baked at 65°C for 4 hours to fully melt paraffin and ensure complete tissue adhesion to the slide surface. Slides were then stored in a desiccator under cold, dry conditions until use. For antigen retrieval, slides were heated to 100°C for 10 minutes, followed by Proteinase K treatment (0.1 μg/ml) for 10-15 minutes, with the exact incubation time held consistent across all samples.

### H&E staining & arthritis severity scores

H&E staining was performed at the University of North Dakota Histology core using standard protocols. To determine arthritis severity scores, a histological analysis of the tibiotarsal joint from each mouse was performed. The sections were evaluated based on arthritis severity on a scale of 0 to 4. Score 0 is given with no inflammation in the tissue. Score 1 with minimal inflammation affecting <5% of tissue. Score 2 with multiple focally extensive areas of inflammation involving 5 % to 25 % of tissue. Score 3 with confluent inflammation involving 25% to 50% of tissue and multiple structures (>3) affected. Score 4 with confluent inflammation in all structures involving >50% of sample [52].

### Spatial transcriptomics with GEOMX

Fresh frozen tibiotarsal joint tissues from both infected and uninfected mice were prepared for the NanoString GeoMX DSP Whole Transcriptome Assay using the manufacturer’s protocol. Joint tissue was immunohistochemically stained with SYTO-13 a blue nucleic acid counter stain (company?), anti-I-E^k^ (company) and anti-Vimentin (company) antibodies.

### Annotation and selection of clusters

Regions of interest (ROI) containing 100 cells each were manually selected from either the later or medial humeroulnar regions in infected mouse joints. Selection was further refined to areas of the synovium in infected mice with morphological markers I-E^k^ and Vimentin being the most concentrated. Anatomically matched ROIs were selected from vehicle-treated control synovium following GeoMx manual slide preparation guidelines.

### Flow Cytometry gating strategy for the analysis of VCAM-1 and ICAM-1

The flow cytometry results were analyzed using FlowJo Software v10.10.1 (Becton, Dickinson and Company). Unstained controls were used to set gates for each fluorochrome, and single-stained controls were used to calculate compensation. The hierarchical gating strategy for the identification of double-positive cells expressing the adhesion molecules CD106 (VCAM-1) and CD54 (ICAM-1) within the CD29^+^/CD90.2^+^ synovial stromal cell population is illustrated with representative plots from control and *Bb* groups at weeks two and four post-infection (see Supplementary Fig S3). At least 100,000 cells were collected per sample. Prior to downstream gating, data quality was assessed using the FlowAI plugin (FlowJo v10.10.1) to automatically detect and quantify anomalous events based on default quality control metrics (flow rate, signal acquisition and dynamic range). Singlets were gated from good events on an FSC-area (FSC-A) versus FSC-H plot to exclude doublets and aggregates. The singlet population was then gated on FSC-A versus side scatter-area (SSC-A) to remove residual debris and select cells for further analysis. Viable cells were selected as cells negative for the viability dye (Fixable Viability Dye APC-Cy7) within the FSC-A/SSC-A gated cells. CD45 expression versus SSC-A on live cells was used to discriminate against hematopoietic lineages (CD45^+^) populations. The CD45^-^ subpopulation was gated for further gating of CD31^-^ events to remove endothelial contaminants. Synovial stromal cells were identified as CD29 (Integrin-b1) and CD90.2 (Thy-1.2) double-positive cells within CD45^-^/CD31^-^ population. Cells double positive for the adhesion molecules CD106 (VCAM-1) and CD54 (ICAM-1) were identified within the CD29^+^/CD90.2^+^ synovial stromal cell gate.

### GeoMx DSP data processing and analysis

Spatial transcriptomic profiling was performed using the NanoString GeoMx® Digital Spatial Profiler (DSP) platform. Following UV-photocleavage of oligo-tagged probes and collection of region-of-interest (ROI/AOI) material, libraries were prepared according to the manufacturer’s protocol and sequenced on an Illumina NovaSeq X. FASTQ files were processed using NanoString GeoMx DSP Analysis Suite software (v2.5.1.166). The final normalized dataset used for downstream analysis was designated *Q3 after filter*.

### Segment-level quality control

Initial quality control (QC) was performed at the AOI (segment) level using the dataset *tech_signal_2*. The following thresholds were applied: ≥1,000 raw reads; ≥80 aligned reads; ≥80 stitched reads; and ≥80 trimmed reads. Sequencing saturation was required to be ≥50 to ensure adequate library complexity. Background signal was assessed using negative control probes (geometric mean ≥2), and No Template Control (NTC) counts were required to be ≤1,000. AOIs were required to have a minimum surface area of 16,000 image-derived units to ensure adequate tissue coverage, and a nuclei count ≥95 to ensure sufficient cellular content. All thresholds were required to be met for AOI retention.

### Probe-level quality control

Probe QC was conducted on the *probe_qc* dataset derived from segment-QC–filtered data. Failed segments were not removed prior to probe QC. Probes were excluded if the ratio of probe geometric mean across all AOIs was <0.1. Probes were also excluded if they failed a Grubbs outlier test in ≥20% of AOIs. Local segment outlier exclusion was not applied. The limit of quantification (LOQ) was defined as 2 standard deviations above background.

### Expression Filtering

Expression filtering was performed at the target level using the dataset *filter_.10*. Targets were retained using the higher of LOQ or user-defined threshold method, in which the minimum expression threshold was defined as the greater of the calculated LOQ or a user-defined count value of 2. Targets were required to meet this threshold in at least 10 AOIs to be retained for downstream analysis.

### Normalization

Normalization was performed using third quartile (Q3) normalization within the GeoMx DSP Analysis Suite. Segment filtering was not applied at this stage; however, target filtering remained enabled. The resulting normalized dataset (*Q3 after filter*) was used for all downstream statistical analyses.

### Differential expression analysis

Differential expression analysis was performed using paired t-tests within the GeoMx DSP Analysis Suite. Three independent paired comparisons were conducted. P-values were adjusted for multiple testing using the Benjamini–Hochberg false discovery rate (FDR) correction. Data processing and visualization were performed using R with the dplyr, ggplot2, ggrepel, and readxl packages. Differential expression data from GeoMX software were imported into R-Studio (Version 2026.01.1+403 (2026.01.1+403)) from a structured Excel file using the *readxl* package, and filtered to remove entries with missing values and resolve gene duplicates. Volcano plots were generated using the *ggplot2* package. The five most significantly up-regulated and five most significantly down-regulated genes were identified by ranking within each group on −log₁₀(p-value) and annotated on the plots. Up-regulated genes were colored red (#D73027), down-regulated genes blue (#2221AC), and non-significant genes grey (#a8a8a8). Threshold lines demarcating the significance and fold-change cutoffs were overlaid in green.

### Spatial transcriptomic data processing and visualization

Downstream analysis was performed in R. Briefly, expression values were extracted from the GeoMx HeatMap Excel export, and regions of interest (ROIs) were assigned to vehicle (uninfected) or *Bb*-infected conditions. Mean expression was computed per condition for each gene, and genes were ranked by the difference in mean expression (infected − vehicle). The top 10 upregulated and top 10 downregulated genes were selected for visualization. Heatmaps were generated using the pheatmap package (v1.0.12), with rows hierarchically clustered using Euclidean distance and complete linkage and columns fixed in condition order (vehicle followed by infected). A diverging color scale (blue–white–red) was anchored at the median expression value across the plotted matrix.

### scRNA-seq differential gene analysis

Pseudobulking analysis of the published scRNA-seq dataset was performed as previously described [53]. Briefly, the synoviocyte subcluster (identified by high expression of *Col1a1* and *Prg4*) was isolated for the differential gene expression analysis. Samples from the same timepoint were split into pseudo-replicates, and the count matrices were generated by summing raw counts for each gene. The differential gene expression at 2 vs. 0 or 4 vs. 0 weeks post-infection was calculated using the DESeq2 package (v.1.50.2) in R.

### GO analysis using EnrichR

Gene ontology (GO) and pathway enrichment analyses were performed using Enrichr (https://maayanlab.cloud/Enrichr), a freely accessible web-based gene set enrichment analysis platform developed by the Ma’ayan Laboratory at the Icahn School of Medicine at Mount Sinai [19]. Separate gene lists of up-regulated and down-regulated differentially expressed genes, as defined by an adjusted p-value threshold of p < 0.05 and an absolute log₂ fold change ≥ 1, were submitted independently to Enrichr for enrichment analysis.

Pathway-level enrichment was assessed against gene set libraries derived from the Molecular Signatures Database (MSigDB) [54]. MSigDB organizes gene sets into collections including hallmark gene sets, which represent well-defined biological states or processes, as well as canonical pathway collections (C2) sourced from resources such as MSigDB, Reactome, and WikiPathways. Enrichment significance was evaluated using a Fisher’s exact test with Benjamini–Hochberg correction for multiple comparisons, and results were ranked by combined score, which integrates the p-value and the deviation from the expected rank.

To investigate associations between differentially expressed genes and metabolic phenotypes, gene lists were additionally queried against metabolite-related gene set libraries available within Enrichr, including those curated from the Metabolomics Workbench and related resources. These libraries link gene sets to known metabolites and metabolic pathways, enabling identification of metabolome-relevant biological processes enriched in the input gene lists. Enrichment was computed and ranked using the same statistical framework applied to the MSigDB pathway analyses.

Bar graph visualizations of enrichment results were generated using the Enrichment Analysis Visualization Appyter, accessed directly from within the Enrichr interface via the “Appyter” [55] to produce bar chart figures ranking the top enriched terms by p-value.

### Immune pathway analysis

Pathway enrichment analysis was performed using Reactome (Build 78 + NCBI_08122021) within the GeoMx DSP Analysis Suite. Pathways were required to meet the following criteria: minimum gene coverage of 20%, minimum of 5 genes detected per pathway, pathway size between 15 and 500 genes, and 10,000 permutations for significance testing. Pathway analysis was conducted separately for each statistical comparison. Immune-related pathways from this list were plotted using ggplot2 [56].

### Venn Diagrams

For each timepoint comparison, genes passing the fold-change threshold were classified by direction (upregulated or downregulated) within each modality. Venn diagrams were constructed to visualise the overlap between the spatial transcriptomics and scRNAseq gene sets, separately for all selected genes, upregulated genes, and downregulated genes. Diagrams were generated using the Shiny App VennDetail. Genes present in the intersection of both modalities and exhibiting the same directional change were classified as ‘concordant’, while genes present in both modalities but with opposing directional changes were classified as ‘discordant’. Genes exceeding the fold-change threshold in only one modality were noted but interpreted with caution, as these may reflect constitutive transcriptional differences between the C3H and C57BL/6 strains rather than genuine biological responses.

### Pathway Analysis

Pathway enrichment analysis was performed using Reactome (Build 78 + NCBI_08122021) within the GeoMx DSP Analysis Suite. Pathways were required to meet the following criteria: minimum gene coverage of 20%, minimum of 5 genes detected per pathway, pathway size between 15 and 500 genes, and 10,000 permutations for significance testing. Pathway analysis was conducted separately for each statistical comparison and plotted using ggplot2 [56].

### Dot plot

To visualize the magnitude and consistency of fold-change across both modalities for the highest-confidence gene sets, dot plots were generated for the top concordant genes at each timepoint comparison, ranked by mean absolute log₂FC across both datasets. For each gene, the log₂FC from the spatial transcriptomics dataset (C3H) and from the scRNAseq synoviocyte dataset (C57BL/6) are shown as individual points, connected by a line to allow direct visual assessment of cross-modality agreement. Genes are displayed on the y-axis and faceted by concordance category (both upregulated or both downregulated). Up to 40 genes are shown per comparison. Dot plots were generated using ggplot2 (4.0.2) in R-Studio. All analyses were performed in R-Studio (Version 2026.01.1+403 (2026.01.1+403)).

## Data availability

Spatial datasets have been deposited in the NCBI Gene Expression Omnibus (GEO) repository and are publicly accessible using accession number GSE327700. Single cell data (published) are also available from NCBI GEO using accession number GSE233850.

## Acknowledgments

We thank Donna Laturnus in the UND histology core for assistance with preparation of slides and H&E staining. We are grateful to Atreyi Ghatak for assistance with mice i.p. injections. We thank the members of the Dhasarathy, Nookala, Petnicki-Ocwieja, and Brissette laboratories, and Dr. Motoki Takaku for helpful discussion, critical evaluation and reading of the manuscript.

## Author contributions

K.G. and Mollie H. performed the wet-lab experiments. V.P. helped with analyses of tissue sections. Y.Z., E.L.A, and T. P-O. helped generate the list of differentially expressed genes from scRNA-seq data. Mike H. and S.J. performed the spatial transcriptomics wet lab and data analyses with GeoMX software, respectively. S.N. helped with flow cytometry and data analyses. C.A.B and A.D. supervised the project, analyzed part of the data, and created the figures. K.G., C.A.B. and A.D. wrote the manuscript.

## Funding and additional information

This work was supported by the Congressionally Directed Medical Research Program (CDMRP)- Tick-borne disease research program (TBDRP) grant [W81XWH-20-1-0168 to A.D. and C.A.B]. Kaitlyn Gura was supported by a National Science Foundation Graduate Research Fellowship (NSF-GRFP) grant number 2334433. Additional support was provided by the National Institute of General Medical Sciences of the National Institutes of Health under Award Numbers U54GM128729, P20GM104360, and P20GM113123.

---

**Supplementary Figure 1.**
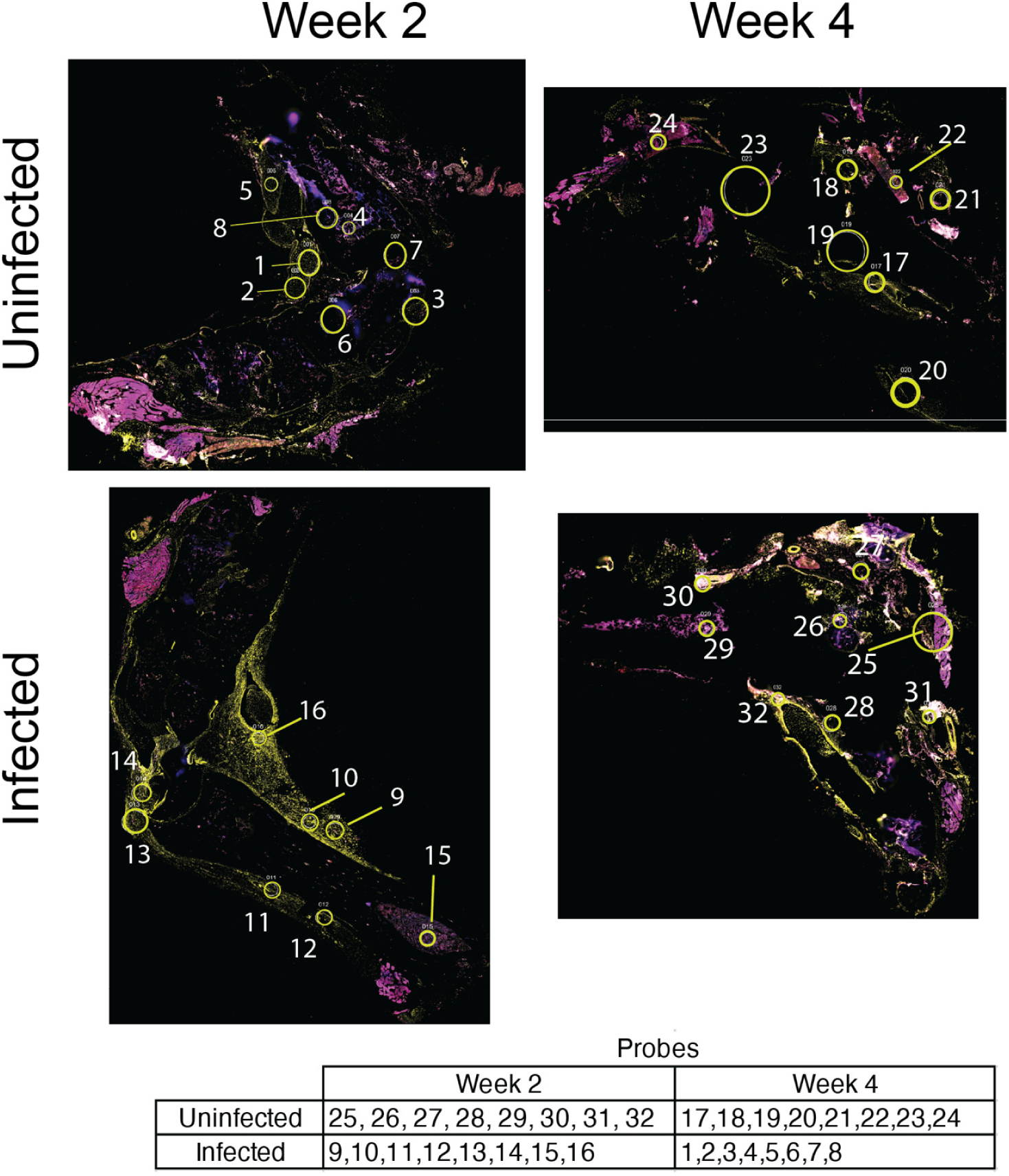
Selection of Regions of Interest and location of probes in *Borrelia burgdorferi*-infected joint tissue. (A) Representative immunofluorescence images of ankle joint sections from uninfected and *Bb*-infected mice at weeks 2 and 4 post-infection. Circles show regions of interest (ROIs) selected for spatial transcriptomics, each circle contained approximated 100 cells. Tissue sections were stained with SYTO-13 (blue; nucleic acid counterstain), anti-I-Eᵏ (red; MHC class II marker), and anti-Vimentin (VIM; yellow). Week 2 Uninfected, Week 2 Infected, Week 4 Uninfected, and Week 4 Infected are shown. (B) Table with probe numbers for each tissue.

**Supplementary Figure 2.**
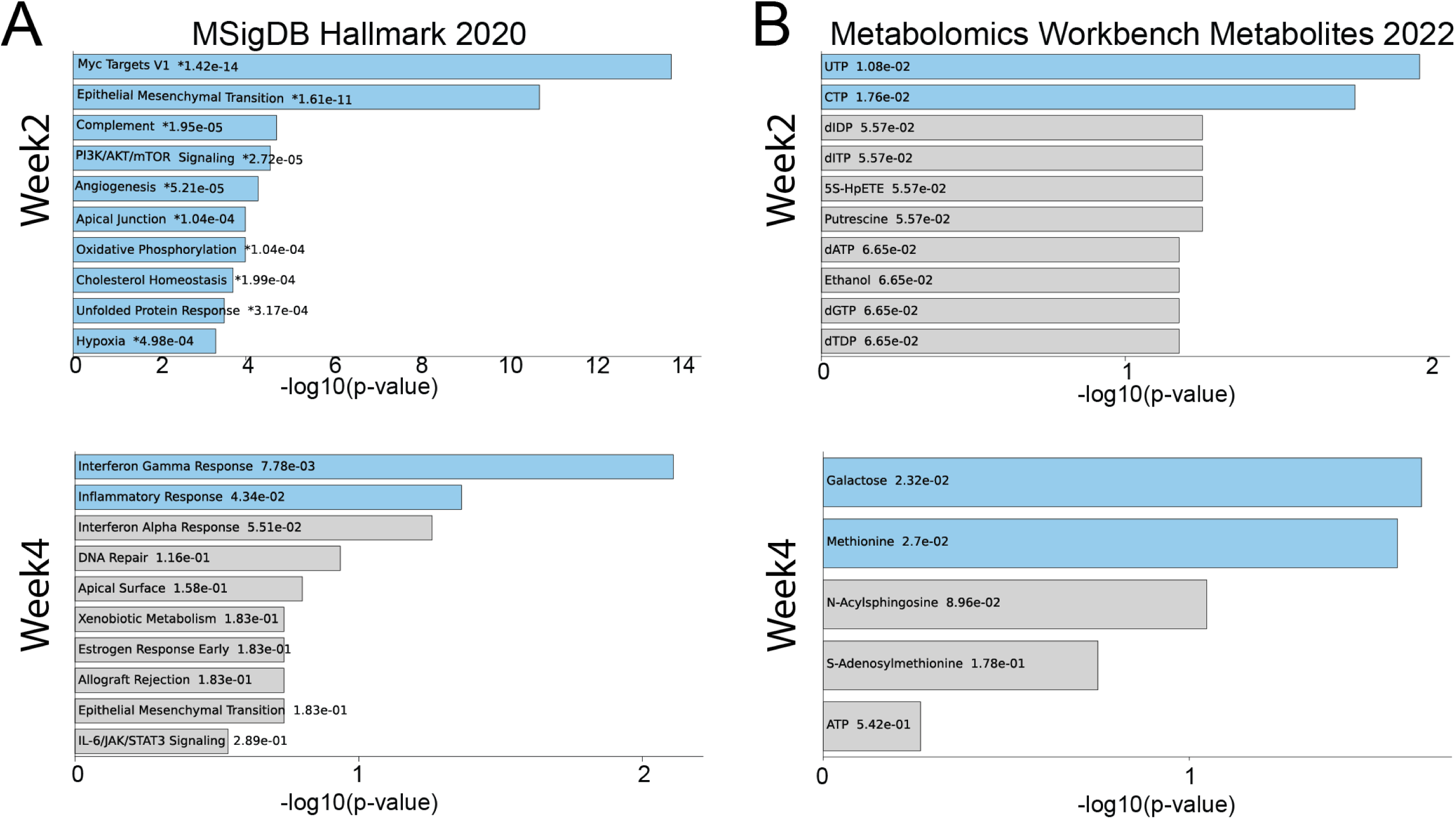
Pathway analysis for downregulated genes following 2 and 4 weeks of *Bb* infection. Top enriched MSigDB Hallmark pathways (A) and Metabolomics metabolites (B) in the downregulated genes at week 2 (top two graphs) and week 4 (bottom two graphs) after *<u>Bb</u>* infection. X axis indicates –log10(p-value) respectively were determined using EnrichR and Appyter. Blue bars are more significant, while gray bars have less significance.

**Supplementary Figure 3.**
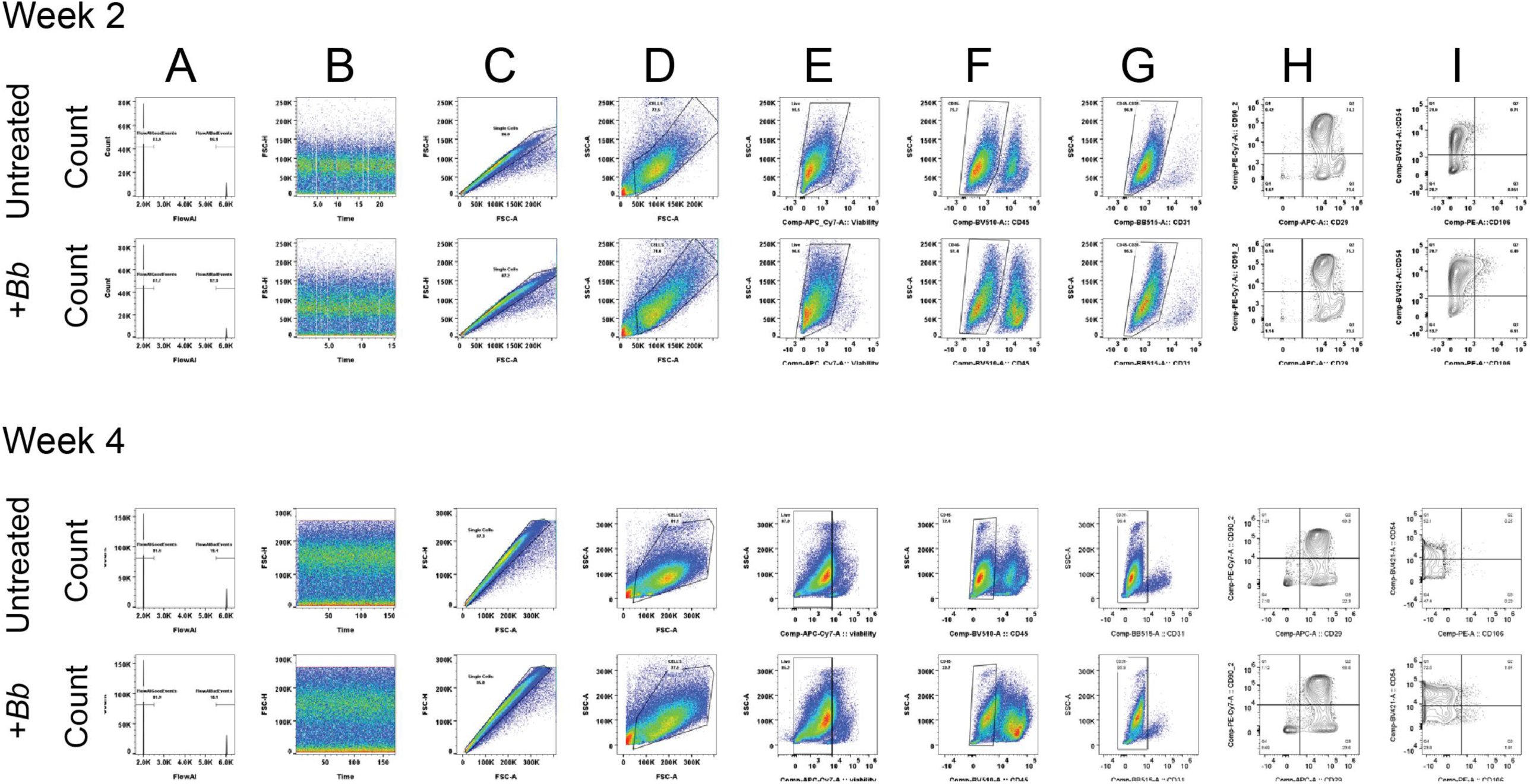
Gating strategy. Unstained controls were used to set gates for each fluorochrome, and single-stained controls were used to calculate compensation. The hierarchical gating strategy for the identification of double-positive cells expressing the adhesion molecules CD106 (VCAM-1) and CD54 (ICAM-1) within the CD29+CD90.2+ synovial stromal cell population is illustrated with representative plots from Vehicle and *Bb* groups at week 2 and week 4 post infection (panels A-I). At least 100,000 were collected per sample. Prior to downstream gating, data quality was assessed using the FlowAI plugin (FlowJo v10.10.1) to automatically detect and quantify anomalous events based on default quality control metrics (flow rate, signal acquisition and dynamic range). A representative histogram (A) and time (x-axis) versus forward scatter-height (FSC-H, y-axis) plot (B) illustrate the distribution of “good events” (retained for further analysis) and bad events (flagged/anomalous) events. Singlets were then gated from good events on an FSC-area (FSC-A) versus FSC-H plot to exclude doublets and aggregates (C). The singlet population was then gated on FSC-A versus side scatter-area (SSC-A) to remove residual debris and select cells for further analysis (D). Viable cells were selected as cells negative for the viability dye (Fixable Viability Dye APC-Cy7) within the FSC-A/SSC-A gated cells (E). CD45 expression versus SSC-A on live cells was used to discriminate against hematopoietic lineages (CD45+) populations (F). The CD45- subpopulation was gated for further gating of CD31- events to remove endothelial contaminants (G). Synovial stromal cells were identified as CD29 (Integrin-b1) and CD90.2 (Thy-1.2) double-positive cells within CD45-CD31- population (H). Cells double positive for the adhesion molecules CD106 (VCAM-1) and CD54 (ICAM-1) were identified within the CD29+CD90.2+ synovial stromal cell gate (I). The flow cytometry results were analyzed using FlowJo Software v10.10.1 (Becton, Dickinson and Company).

**Supplementary Figure 4.**
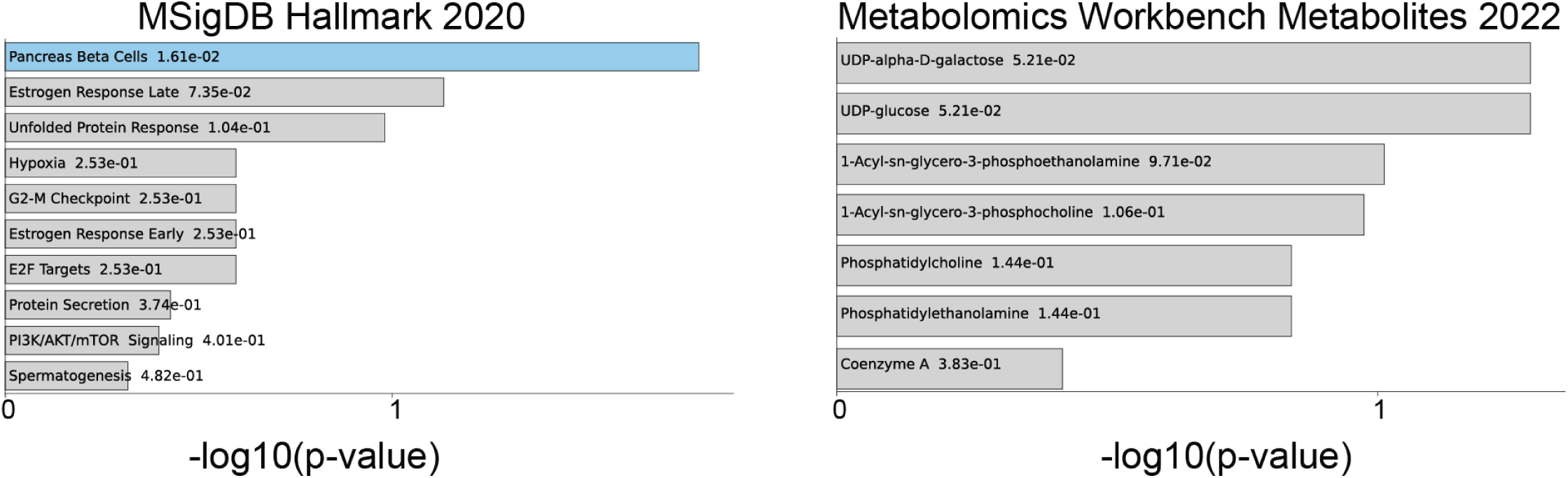
**Pathway analysis for upregulated genes in lateral synovium after 2 weeks of Bb infection**. Top enriched MSigDB Hallmark pathways (A) and Metabolomics metabolites (B) in the upregulated genes at week 2 after Borrelia infection. X axis indicates –log10(p-value) respectively were determined using EnrichR and Appyter. Blue bars are more significant, while gray bars have less significance.

